# Hepatocyte Growth Factor/MET Activator Rescues Working Memory Deficits After Repeated Mild Traumatic Brain Injury

**DOI:** 10.1101/2025.09.25.678537

**Authors:** Katelyn A. Martino, Arush Nakhre, Renee M. Demarest, David M. Devilbiss

## Abstract

Mild traumatic brain injury (**mTBI**) can produce persistent cognitive and behavioral deficits. These impairments result in a reduced quality of life and difficulty returning to work, school, or other activities. Individuals with repeated injuries show increased risk for greater cognitive impairment and persistence of symptoms. A variety of pharmacological approaches have been tried to limit cognitive symptoms and other aspects of secondary injury following mTBI. However, their efficacy and ability to treat the sequela of mTBI remains disputed and no FDA-approved drug exists for mTBI. However, neurotrophins have considerable promise as regenerative therapies for mTBI by exhibiting procognitive, neuroprotective, and anti-inflammatory actions. One neurotrophin, hepatocyte growth factor and its receptor MET (HGF/MET) are upregulated in response to CNS injury within the prefrontal cortex and other regions supporting memory and higher cognitive function impaired by TBI. HGF/MET activation can be anti-inflammatory and neuroprotective, yet an understanding of its actions on cognitive function after mTBI is limited. Using a closed-head midline impact model of mild TBI, we characterized the actions of the HGF/MET system on working memory performance after repeated injury. Following repeated mild TBI, the HGF/MET positive modulator dose-dependently rescued the working memory deficits following injury. These actions indicate that peptidergic transmitter systems including HGF/MET may hold critical pharmacological targets for treating the neurosequela of TBI.

## Introduction

Mild traumatic brain injury (**mTBI**) is a serious health concern affecting nearly 30-50 million people/year worldwide^1-3^. Impaired working memory, attention, and executive function are common symptoms of TBI and are significant predictors of long-term productivity and quality of life after injury^4,5^. Working memory is critical for sustained attention, goal-directed behaviors, and cognitive flexibility affected by mTBI. Specifically, working memory is the temporary and active maintenance of information when it is no longer perceptually present for updating goal/task-directed behaviors^6,7^. These executive processes are dependent on the prefrontal cortex (**PFC**) of rodents and humans, an area that is substantially vulnerable to TBI^8,9^. Repetitive mTBI can produce more severe, longer-lasting cognitive impairments and neuropathology than a single injury and can result in outcomes similar to severe injury^10-13^. Repetitive mTBI increases the risk for neurodegenerative diseases and accelerates the accumulation of pathological hallmarks in the brain^14^. Given that the incident rate for repeated mTBI is 5.6–36% across the general population^15^, developing effective cognitive treatments remains an unmet need and critical gap in the field of traumatic brain injury^16^.

There are several classes of growth factors that act within the brain by binding to tyrosine kinase receptors including hepatocyte growth factor (**HGF**) and prototypical neurotrophins such as nerve growth factor and brain-derived growth factor^17^. Each growth factor and receptor can activate a distinct and overlapping array of signaling pathways in selective cell populations leading to modulation of neurotransmission and synaptic plasticity. Within the brain, activation of HGF signaling and its receptor MET can be strongly neuroprotective^18,19^, promote neurogenesis^19-21^, and enhance spinogenisis^22^. HGF and MET are upregulated after brain injury and have been extensively studied in animal models of spinal cord injury^23^. Moreover, MET is highly expressed in cortical areas involved in attention, memory, and sensorimotor functions^24^. Studies in humans and rodent models of neurodegenerative diseases demonstrate that activating HGF/MET signaling results in cognitive enhancement likely through the potentiation of NMDA currents and increased dendritic arborization^16,25,26^. However, the effect of HGF/MET on recovery from mTBI and the role of activating the HGF/MET signaling in working memory remains unclear. In these studies we used dihexa, a first-generation positive modulator of MET by potentiating the dimerization of HGF^27^, to determine the degree to which HGF/MET activation affects PFC-dependent spatial working memory in rats. Combined these data provide novel insight into the therapeutic actions of HGF/MET activation to ameliorate cognitive deficits after repeated mTBI.

## Materials and Methods

Detailed Methods and Materials are provided as a Supplementary File.

### Animals and Surgery

Male (n=33) and female (n=19) Long-Evans rats were single-housed with enrichment on a 12/12-hour reverse light cycle and maintained at 85–100% free-feeding weight (5% bodyweight of chow/day). A reverse light cycle ensured that repeated sham or closed-head mTBI surgeries (n=3 over 1 week) and training/testing the working memory task^28-30^ occurred during their active period (**Figure 1A**). For surgery, the midline of the skull was exposed while animals were under isoflurane anesthesia (2–3% in 95%O2:5%CO2). The animal was removed from anesthesia and placed on a Marmarou foam block^31,32^ and impacted midline (-2.5mm AP; 5.5m/s, 2.5mm depth, 100ms dwell, 5mm hemispherical tip). After measuring the righting reflex, animals were returned to anesthesia to close the incision. Resting prone on the foam block allowed the animal’s head to accelerate and move unrestrained following impact. Cannula were placed in the lateral ventricle (26ga.; unilateral right side; -1.0AP,-1.4ML,-1.4DV) after re-anesthetization during the third surgery. Sham animals experienced identical surgical procedures but did not receive mTBI. All procedures were in accordance with NIH guidelines and approved by the Rowan University Institutional Animal Care and use Committee.

**Figure 1.**
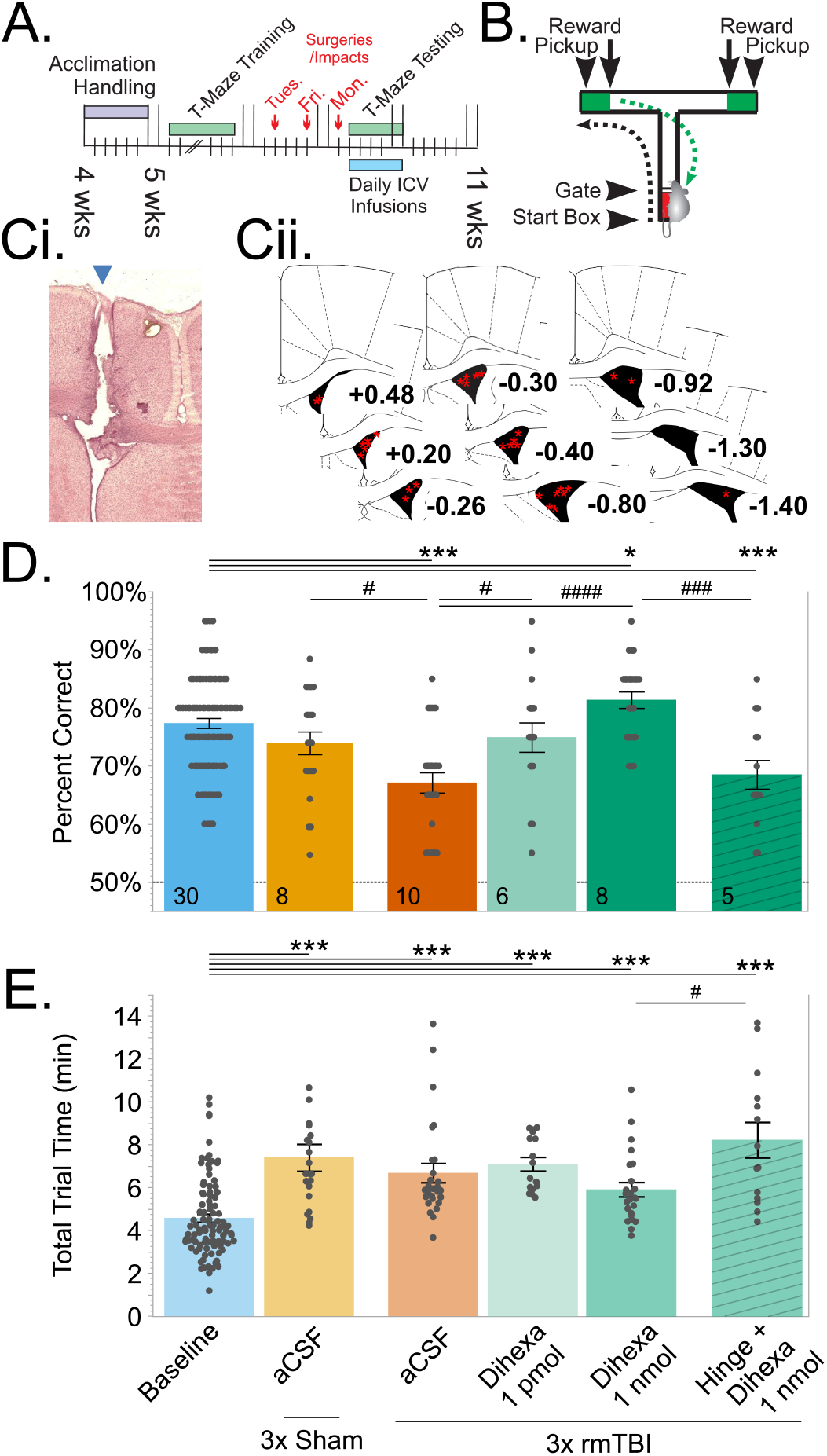
Intracerebroventricular (ICV) infusion of the HGF/MET positive modulator, dihexa rescues working memory performance after repeated mTBI. A) Experimental timeline illustrating repeated injuries, drug administration, and T-maze testing. B) During T-maze testing, the rat is released from the start box after a delay period, receives a hand-fed reward for a correct response after choosing the arm not entered in the prior trial (black arrow), and removal of the animal from the maze (green arrow) to begin another trial. C) Histological localization of infusion needle placement. Ci) Photomicrograph of an exemplar neutral red stained coronal section illustrating the cannula/infusion needle tract entering the lateral ventricle (blue arrow; 0.0 AP). Image represents a stitched series of 20x images. Cii) Anatomical drawing series illustrates the final placement of cannula for each animal. Animals with infusion needles not within the lateral ventricle were excluded from analyses. D) Performance in the T-maze delayed alternation task of working memory during Baseline (pre injury; 3 day mean) and after repeated mTBI 3rd - 5th days of treatment. 20 trials; 50% is chance performance. E) Total duration of T-maze testing for animals in (D). The highest dose of dihexa tested (1 nmol ICV) improves performance and trial times in repetitive mTBI animals. MET antagonists block dihexa’s effects. N’s indicated in bar insets. *p<0.05, **p<0.01, Dunnett’s; #p<0.05, ##p<0.01, ###p<0.001, Tukey’s HSD. T-maze testing was performed 5 min after ICV infusions each day.

### Spatial working memory testing

Animals were trained in a discrete-trial, rewarded T-maze delayed-nonmatch to position task (Fig. 1B)^29,33-36^ to achieve stable performance of 80±5% correct over 3 consecutive days (20 trials; 1 session/day; 60db white noise). If needed, delays were increased in 5sec increments during training (max 15sec) to maintain performance at criterion while waiting for a surgical window. Once criterion was met, animals were provided *ad lib* food and underwent repeated sham or mTBI surgeries, returned to food regulation, assigned an inter-trial delay 5-sec. longer than baseline, and tested daily in the T-maze task (days 1-5 post final surgery; Fig. 1A). Testing sessions were conducted identically to training sessions.

### Drug Administration

Dihexa (MedChemExpress Inc.; 1.0pmol or 1.0nmol/2uL) was dissolved in 2.5% dimethyl sulfoxide (**DMSO**) and Dulbecco’s Phosphate Buffered Saline (**DPBS**;VWR #02–0117). The peptide MET antagonist, Hinge (KDYIRN, CHI Scientific; 300pmol/2μL) was dissolved in DPBS. An intracerebroventricular (**ICV**) infusion of DPBS or dihexa (30ga needle; RWD Life Sciences Inc.) was made daily into awake, unrestrained animals 5 minutes before behavioral testing with infusion needles remained in the guide cannula for 30 seconds after infusion. Administration of Hinge occurred 5 minutes before the dihexa infusion.

### Histology

Following T-maze testing, animals were anesthetized (isoflurane; 5% in 95%O_2_:5%CO_2_) and perfused with 0.9% saline followed by 10% formalin. Methylene Blue (1% in DPBS; 2-5μL) was injected through the ICV infusion needle. Brains were extracted and stored in 50mL of 10% formalin (>24hrs) followed by 30% sucrose (>48hrs). Frozen 40μm coronal sections were collected and counterstained with Neutral Red to visualize the final surgical placement of the infusion needle within the lateral ventricle.

#### Scatter Assay

To confirm dihexa augments HGF/MET signaling pathways, Madin-Darby canine kidney (MDCK;ATCC #CCL-34) cells were grown in six-well plates to form isolated, confluent colonies. Colonies were washed twice with DPBS and media was replaced with serum-free Dulbecco’s modified Eagle’s medium. DPBS, dihexa (10-1000 pM), HGF (0.34μM), or dihexa (1000pM)+HGF (0.34μM) was added to the media and plates were incubated at 37°C with 5% CO_2_ for 10 hours. Media was removed and cells were fixed with 3.7% formalin in phosphate buffered saline and immunofluorescent labeling performed for E-Cadherin (#3195T;Cell Signaling Technology Inc.).

#### Quantification and Statistical Analysis

All analyses were performed within the mixed model framework of JMP (ver.18.2.1, JMP Statistical Discovery). Tukey’s HSD post-hoc test was used to evaluate the effect of Injury, Surgery#, and Sex on animal weights and righting reflex times. Tukey’s HSD and Dunnett-Hsu post-hoc test were used to compare performance in the T-maze task. E-Cadherin labeling of MDCK cells was quantified by imaging cell cultures (Keyence BZ-X710) and identifying cell membranes with Cellpose-SAM^37^. Identified membranes were scaled by 80% to quantify E-Cadherin labeling intensity within the cytoplasm. A mixed model was used with Tukey’s HSD and Dunnett-Hsu to compare drug effects.

## Results

### Immediate physiological effects of repeated mild TBI

For the first of three repeated surgeries, no differences in weights were found between injury groups ANOVA_(sexXinjury)_ F_(1,48)_= 0.5856, p=0.4479). Males (204.3g; range 160-296g) initially did weigh more than females (183.6g; range 150-233; 111% heavier; ANOVA_(sex)_ F_(1,48)_= 5.5916, p=0.0221. Animal weight increased in a sex-dependent manner during the series of repeated surgeries (ANOVA_(sexXsurgery)_ F_(2,96)_= 20.5374, p<0.0001; Supplemental Table 1) and was not affected by the type of injury (ANOVA_(sexXsurgeryXinjury)_ F_(2,96)_= 0.0592, p=0.9425). Apnea after injury was not observed, and there were no mortalities. Over the sequence of repeated surgeries, righting reflex times were significantly longer after mild TBI (ANOVA(injury) F(1,48) =25.1215, p<0.0001, 1^st^ surgery: 4:15.4±24.8 vs mTBI: 6:42.7±29.5 mean±SEM, mm:ss.0) and decreased with each additional injury (ANOVA (surgery) F(2,96) =9.4537, p=0.0002, Supplemental Table 2; 2^nd^ surgery: 2:23.8±19.2 vs mTBI: 5:18.2±10.5, 3^rd^ surgery: 2:41.7±17.7 vs mTBI: 4:24.6±31.6). No sex-dependent differences were found (ANOVA_(sexXinjury)_ F_(1,48)_= 0.0154, p=0.9017) and there was no interaction between righting reflexes during each repeated surgery and type of injury (sham/mTBI; ANOVA_(surgeryXinjury)_ F_(2,96)_=0.9115, p=0.4054).

### HGF/MET activation rescues working memory performance after repetitive mild TBI

Baseline performance on the working memory task was 77.4 ±1% correct with a mean delay of 5sec (range 5-15 sec) for all animal groups. Repetitive mild TBI significantly impaired performance from baseline (Fig. 1. ANOVA_(injury)_ F_(2,136)_=14.4199, p<0.0001; HSD, p<0.0001, 67.1% correct) and from sham injured animal performance (HSD, p=0.0159). Repetitive sham did not impair working memory performance (HSD; p=0.6592, 75.0% correct). No sex-dependent differences were found (ANOVA_(sex)_ F_(1,30)_=3.0877, p=0.0888; Supplemental Figure 1A,B) and were combined for all analyses. Comparing animal groups that received repeated mTBI (ANOVA F_(4,170)_=15.5705, p<0.0001), subchronic administration of dihexa (days 1-5) dose-dependently rescued working memory performance (days 3-5; Fig. 1D). Following the highest dose of dihexa (1nmol/day ICV), working memory performance in repetitively injured animals was significantly higher than those receiving vehicle (81.5% vs. 67.1%, HSD p<0.0001) and animal performance during baseline (77.4% vs. 77.6%, Dunnett’s; p=0.0258). Pretreatment with the MET antagonist, Hinge (KYRDIN, 300pmol) significantly blocked the procognitive effects of dihexa (1 nmol/day, 68.6%, HSD p<0.0001). Post-surgical performance was assessed on days 3-5, representing stable performance after animals re-equilibrated after reinstatement of food regulation.

The T-maze working memory task was completed within 4:38(min:sec)±11sec (21 trials total, mean±SEM) under baseline conditions. Total session length was significantly increased following both repetitive sham and mild TBI surgeries (Fig. 1E., ANOVA F_(2,131)_=27.0651, p<0.0001), totaling 7:28±38sec (Dunnett’s p<0.0001) and 6:445±29sec (Dunnett’s p<0.0001) after sham or mild TBI respectively. No sex-dependent differences were found (ANOVA_(sex)_ F_(1,30)_= 0.0241, p=0.8777, Supplemental Figure 1C,D) and were again combined for all analyses. Dihexa (1.0nmol ICV) did limit increased total session time compared to saline treatment after repeated mTBI (5:58 ±20 sec), although this effect was not statistically significant (HSD, p=0.9777). However, total session time following pretreatment with Hinge (KYRDIN, 300 pmol) was significantly greater than baseline (8:17±50; Dunnett’s p<0.001) and was significantly greater than dihexa treated animals (1.0nmol ICV; HSD, p=0.0161).

### HGF/MET signaling is enhanced following dihexa administration in a scatter-assay

To confirm dihexa augments HGF/MET signaling pathways, internalization of E-cadherin was examined in the MDCK cell line (Figure 2). E-cadherin is redistributed from the cell membrane into the cytoplasm in order to promote cell migration^38^. Analysis of fluorescence intensity within the cytoplasm (ANOVA F_(5,141)_=21.0631, p<0.0001), demonstrated that E-cadherin was internalized after application of HGF (0.34μM; 10.3±0.6 mean pixel value; HSD, p<0.0001); whereas under vehicle control (DPBS, 6.2±0.5) conditions, labeling of E-cadherin was primarily membrane-bound as expected (6.2±0.5 mean pixel value). Dihexa treatment alone (Fig 2C), dose-dependently increased E-cadherin internalization from 10 pM (6.3 ±0.2) to the highest dose of tested (100pM, 6.6±0.3; 1.0nM, 8.6±0.4 mean pixel value, HSD p=0.0007). The combination of dihexa and HGF essentially eliminated labeling of membrane-bound E-cadherin (10.3±0.5; HSD, p<0.0001). Together these data support the proposed mechanism that dihexa facilitates dimerization of HGF that subsequently activates the MET receptor and, subsequently, downstream intracellular signaling pathways^22^.

**Figure 2.**
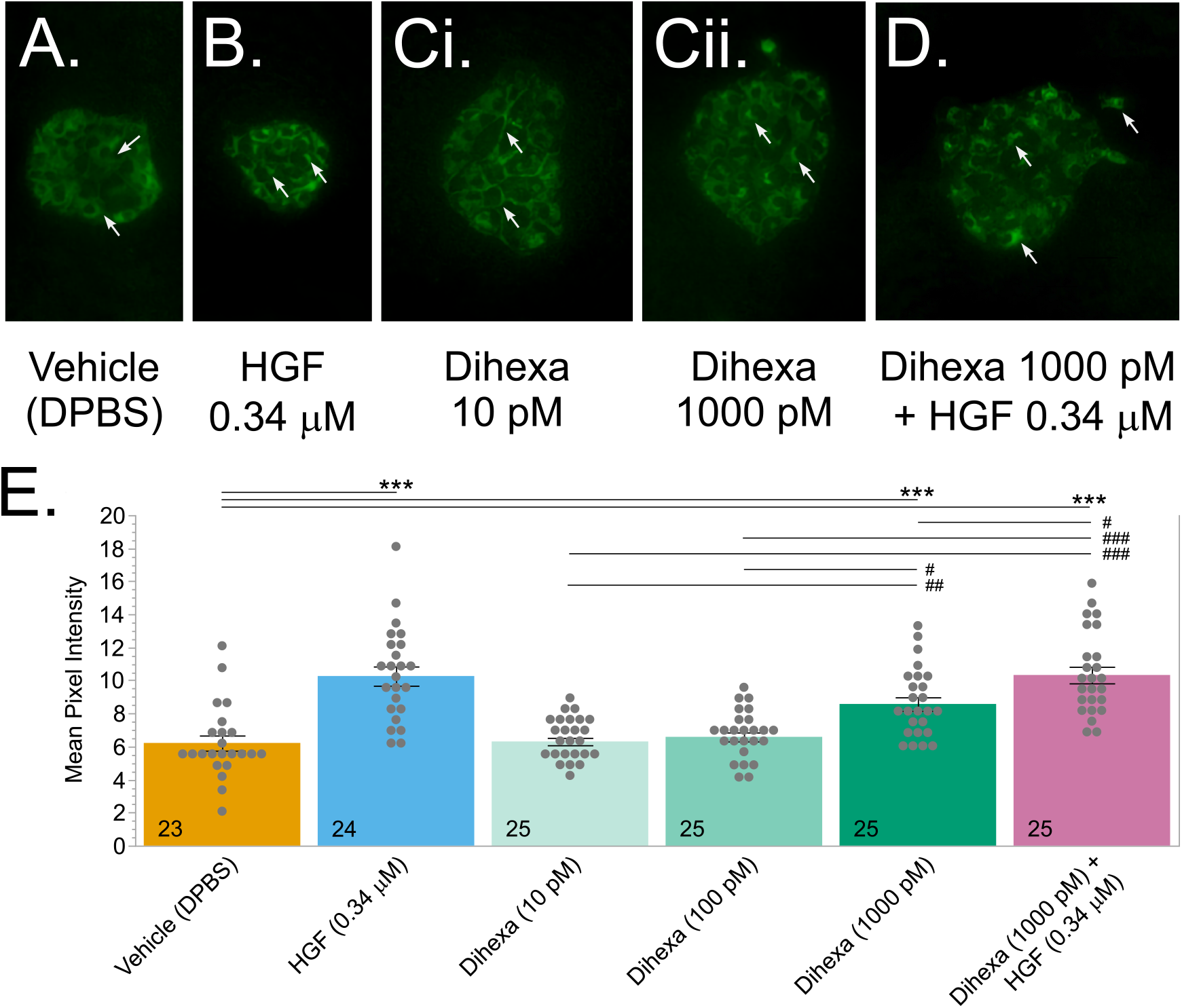
E-cadherin markers of cell-cell adhesion following MET activation in representative MOCK cell colonies. A) Colonies treated with Vehicle (Dulbecco’s Phosphate Buffered Saline, DPBS) demonstrate diffuse E-cadherin (Green) labeling on cell membranes. B) Hepatocyte growth factor (HGF) activation of MET results in migration of E-cadherin to the cytosol immediately prior to scattering. C) Dihexa treatment alone, dose-dependently increases internalization of E-cadherin to the cytosol. D) Dihexa facilitates the actions of HGF to internalize E-cadherin to the cytosol. All images are taken at 40x. E) Quantification of E cadherin labeling in the cytoplasm for a random sampling of cells across multiple colonies. Mean pixel intensity (lower values= less label). N’s indicated in bar insets. *p<0.05, **p<0.01, Dunnett’s; #p<0.05, ##p<0.01, ###p<0.001, Tukey’s HSD.

## Discussion

Treatment of TBI with neurotrophic factors holds substantial promise and this study represents the first evidence that a small molecule activator of the HGF/MET system can improve cognitive function following repeated mild TBI. In this study, we found dihexa ameliorated deficits in working memory performance in rats following a repetitive mild TBI. We further observed that the procognitive effects of dihexa were blocked by Hinge, an HGF/MET dimerization antagonist, indicating these actions are mediated by HGF/MET signaling pathways.

This is the first evidence that activation of HGF/MET signaling directly improves recovery of cognitive function after TBI, the effects of HGF treatment in preclinical models of neurodegenerative models^17,27,39,40^ and spinal cord injury (SCI)^41,42^ have been well studied. Following SCI and cerebral ischemic injury, HGF and MET are upregulated after injury suggesting a role in the neuroprotective response after insult^43^, but see^44^. The mechanisms supporting the procognitive actions of MET are poorly understood, though NMDAR/Ca^2+^ are posited to underlie these actions^22,25,40^. Indeed, we were able to confirm that dihexa can increase Ca^2+^-dependent cellular processes as endocytosis of E-cadherin is dependent on elevated intracellular Ca^2+^ signaling following MET activation^38^. Addressing the complement of signaling pathways activated by MET casually associated with its prognitive actions will be important for understanding the role of HGF and other growth factors in cognitive function.

## Conclusions

The current observations indicate the HGF/MET activator, dihexa can dose-dependently ameliorate working memory deficits in a rodent model of repeated mild TBI. This action may be dependent on increased intracellular Ca^2+^ signaling following MET activation. Together, these findings indicate that HGF and possibly other growth factors hold promise for treating the neurocognitive sequela following TBI as small molecule HGF/MET activators currently in clinical trials may provide an accelerated path for testing its clinical efficacy in TBI patient populations^40^.

### Transparency, Rigor, and Reproducibility Statement

The goal of this study was to characterize the effects of MET activation on working memory performance after repeated mild TBI. Surgeries were performed by experienced researchers and behavioral testing was performed blind of injury status and therapeutic interventions. Synthesized lots of dihexa were validated and assayed for impurities using mass spectrometry. Manufacturing lots were found to be indistinguishable. Statistical analyses were conducted using JMP Pro 18.0.2 and included Mixed Model ANOVA with Tukey’s HSD post-hoc comparisons. Data from this study are available in a FAIR data repository (odc-tbi.org). The study design and analytic plan were not preregistered.

## Acknowledgments

The authors would like to thank Doug Fox, Michael Ingling, Joseph Kalish, Maurice Linder-Jackson, Sarah Marcum, Laura Milovic, Arwa Muhamed, and Ashley Thayaparan for their technical assistance in the working memory task. The authors would also like to thank Benrd Spur, Ana Rodriguez Rosich, Mikhail Anikin for their MALDI-TOF and mass spectrometry and assistance characterizing synthesized dihexa.

## Authorship confirmation/contribution statement

K Martino: Writing – Original Draft, Data Curation, Investigation, Visualization

A Nakhre: Writing – Original Draft, Investigation

RM Demarest: Conceptualization, Resources, Supervision (MDCK scatter assay)

DM Devilbiss – Conceptualization, Formal Analysis, Funding Acquisition, Methodology, Resources, Supervision, Validation, Writing – Review & Editing

## References

1. Faul M, Xu LZ, Wald MW, et al. Traumatic brain injury in the United States: emergency department visits, hospitalizations and deaths 2002–2006. Centers for Disease Control and Prevention, National Center for Injury Prevention and Control: Atlanta (GA); 2010.

2. Bruns J, Jr., Hauser WA. The epidemiology of traumatic brain injury: a review. Epilepsia 2003;44(10):2–10, doi:10.1046/j.1528-1157.44.s10.3.x

3. Maas AIR, Menon DK, Adelson PD, et al. Traumatic brain injury: integrated approaches to improve prevention, clinical care, and research. Lancet Neurol 2017;16(12):987–1048, doi:10.1016/S1474-4422(17)30371-X

4. Wood RL, Worthington A. Neurobehavioral Abnormalities Associated with Executive Dysfunction after Traumatic Brain Injury. Frontiers in behavioral neuroscience 2017;11(195, doi:10.3389/fnbeh.2017.00195

5. Wood RL, Rutterford NA. Demographic and cognitive predictors of long-term psychosocial outcome following traumatic brain injury. J Int Neuropsychol Soc 2006;12(3):350–8, doi:10.1017/s1355617706060498

6. Baddeley AD, Hitch GJ. Developments in the concept of working memory. Neuropsychology 1994;8(4):485–493, doi:10.1037/0894-4105.8.4.485

7. Smith EE, Jonides J. Storage and executive processes in the frontal lobes. Science 1999;283(5408):1657–61, doi:10.1126/science.283.5408.1657

8. McAllister TW. Neurobiological consequences of traumatic brain injury. Dialogues Clin Neurosci 2011;13(3):287–300

9. Smith CJ, Xiong G, Elkind JA, et al. Brain Injury Impairs Working Memory and Prefrontal Circuit Function. Front Neurol 2015;6(240, doi:10.3389/fneur.2015.00240

10. Rubin TG, Lipton ML. Sex Differences in Animal Models of Traumatic Brain Injury. J Exp Neurosci 2019;13(1179069519844020, doi:10.1177/1179069519844020

11. Karton C, Blaine Hoshizaki T. Concussive and subconcussive brain trauma: the complexity of impact biomechanics and injury risk in contact sport. Handb Clin Neurol 2018;158(39-49, doi:10.1016/B978-0-444-63954-7.00005-7

12. Guskiewicz KM, McCrea M, Marshall SW, et al. Cumulative effects associated with recurrent concussion in collegiate football players: the NCAA Concussion Study. JAMA 2003;290(19):2549–55, doi:10.1001/jama.290.19.2549

13. McCrory P, Collie A, Anderson V, et al. Can we manage sport related concussion in children the same as in adults? Br J Sports Med 2004;38(5):516–9, doi:10.1136/bjsm.2004.014811

14. Uryu K, Laurer H, McIntosh T, et al. Repetitive mild brain trauma accelerates Abeta deposition, lipid peroxidation, and cognitive impairment in a transgenic mouse model of Alzheimer amyloidosis. J Neurosci 2002;22(2):446–54

15. Greco T, Ferguson L, Giza C, et al. Mechanisms underlying vulnerabilities after repeat mild traumatic brain injuries. Exp Neurol 2019;317(206-213, doi:10.1016/j.expneurol.2019.01.012

16. Leddy JJ, Baker JG, Willer B. Active Rehabilitation of Concussion and Post-concussion Syndrome. Phys Med Rehabil Clin N Am 2016;27(2):437–54, doi:10.1016/j.pmr.2015.12.003

17. Atkinson E, Dickman R. Growth factors and their peptide mimetics for treatment of traumatic brain injury. Bioorg Med Chem 2023;90(117368, doi:10.1016/j.bmc.2023.117368

18. Doeppner TR, Kaltwasser B, ElAli A, et al. Acute hepatocyte growth factor treatment induces long-term neuroprotection and stroke recovery via mechanisms involving neural precursor cell proliferation and differentiation. J Cereb Blood Flow Metab 2011;31(5):1251–62, doi:10.1038/jcbfm.2010.211

19. Shang J, Deguchi K, Yamashita T, et al. Antiapoptotic and antiautophagic effects of glial cell line-derived neurotrophic factor and hepatocyte growth factor after transient middle cerebral artery occlusion in rats. J Neurosci Res 2010;88(10):2197–206, doi:10.1002/jnr.22373

20. Nicoleau C, Benzakour O, Agasse F, et al. Endogenous hepatocyte growth factor is a niche signal for subventricular zone neural stem cell amplification and self-renewal. Stem Cells 2009;27(2):408–19, doi:10.1634/stemcells.2008-0226

21. Wang M, Gamo NJ, Yang Y, et al. Neuronal basis of age-related working memory decline. Nature 2011;476(7359):210–3, doi:10.1038/nature10243

22. Benoist CC, Kawas LH, Zhu M, et al. The procognitive and synaptogenic effects of angiotensin IV-derived peptides are dependent on activation of the hepatocyte growth factor/c-met system. The Journal of pharmacology and experimental therapeutics 2014;351(2):390–402, doi:10.1124/jpet.114.218735

23. Kitamura K, Nagoshi N, Tsuji O, et al. Application of Hepatocyte Growth Factor for Acute Spinal Cord Injury: The Road from Basic Studies to Human Treatment. Int J Mol Sci 2019;20(5), doi:10.3390/ijms20051054

24. Thewke DP, Seeds NW. The expression of mRNAs for hepatocyte growth factor/scatter factor, its receptor c-met, and one of its activators tissue-type plasminogen activator show a systematic relationship in the developing and adult cerebral cortex and hippocampus. Brain Res 1999;821(2):356–67, doi:10.1016/s0006-8993(99)01115-4

25. Kato T, Funakoshi H, Kadoyama K, et al. Hepatocyte growth factor overexpression in the nervous system enhances learning and memory performance in mice. J Neurosci Res 2012;90(9):1743–55, doi:10.1002/jnr.23065

26. Akimoto M, Baba A, Ikeda-Matsuo Y, et al. Hepatocyte growth factor as an enhancer of nmda currents and synaptic plasticity in the hippocampus. Neuroscience 2004;128(1):155–62, doi:10.1016/j.neuroscience.2004.06.031

27. Wright JW, Harding JW. The Brain Hepatocyte Growth Factor/c-Met Receptor System: A New Target for the Treatment of Alzheimer’s Disease. J Alzheimers Dis 2015;45(4):985–1000, doi:10.3233/JAD-142814

28. Devilbiss DM, Spencer RC, Berridge CW. Stress Degrades Prefrontal Cortex Neuronal Coding of Goal-Directed Behavior. Cereb Cortex 2016, doi:10.1093/cercor/bhw140

29. Berridge CW, Devilbiss DM, Andrzejewski ME, et al. Methylphenidate preferentially increases catecholamine neurotransmission within the prefrontal cortex at low doses that enhance cognitive function. Biol Psychiatry 2006;60(10):1111–1120, doi:10.1016/j.biopsych.2006.04.022

30. Horvat L, Foschini A, Grinias JP, et al. Repetitive mild traumatic brain injury impairs norepinephrine system function and psychostimulant responsivity. Brain Res 2024;1839(149040, doi:10.1016/j.brainres.2024.149040

31. Marmarou CR, Prieto R, Taya K, et al. Marmarou Weight Drop Injury Model. In: Animal Models of Acute Neurological Injuries. (Chen J, Xu ZC, Xu X-M, et al. eds.) Humana Press: Totowa, NJ; 2009; pp. 393–407.

32. Marmarou A, Foda MA, van den Brink W, et al. A new model of diffuse brain injury in rats. Part I: Pathophysiology and biomechanics. J Neurosurg 1994;80(2):291–300, doi:10.3171/jns.1994.80.2.0291

33. Devilbiss DM, Berridge CW. Cognition-enhancing doses of methylphenidate preferentially increase prefrontal cortex neuronal responsiveness. BiolPsychiatry 2008;64(7):626–635

34. Devilbiss DM, Jenison RL, Berridge CW. Stress-induced impairment of a working memory task: role of spiking rate and spiking history predicted discharge. PLoS Comput Biol 2012;8(9):e1002681, doi:10.1371/journal.pcbi.1002681

35. Spencer RC, Devilbiss DM, Berridge CW. The cognition-enhancing effects of psychostimulants involve direct action in the prefrontal cortex. Biol Psychiatry 2015;77(11):940–50, doi:10.1016/j.biopsych.2014.09.013

36. Berridge CW, Devilbiss DM, Martin AJ, et al. Stress degrades working memory-related frontostriatal circuit function. Cereb Cortex 2023;33(12):7857–7869, doi:10.1093/cercor/bhad084

37. Pachitariu M, Rariden M, Stringer C. Cellpose-SAM: superhuman generalization for cellular segmentation. bioRxiv 2025;2025.04.28.651001, doi:10.1101/2025.04.28.651001

38. Delva E, Kowalczyk AP. Regulation of cadherin trafficking. Traffic 2009;10(3):259–67, doi:10.1111/j.1600-0854.2008.00862.x

39. Sun X, Deng Y, Fu X, et al. AngIV-Analog Dihexa Rescues Cognitive Impairment and Recovers Memory in the APP/PS1 Mouse via the PI3K/AKT Signaling Pathway. Brain Sci 2021;11(11), doi:10.3390/brainsci11111487

40. Moebius HJ, Church KJ. The Case for a Novel Therapeutic Approach to Dementia: Small Molecule Hepatocyte Growth Factor (HGF/MET) Positive Modulators. J Alzheimers Dis 2023;92(1):1–12, doi:10.3233/JAD-220871

41. Desole C, Gallo S, Vitacolonna A, et al. HGF and MET: From Brain Development to Neurological Disorders. Front Cell Dev Biol 2021;9(683609, doi:10.3389/fcell.2021.683609

42. Sakai K, Aoki S, Matsumoto K. Hepatocyte growth factor and Met in drug discovery. J Biochem 2015;157(5):271–84, doi:10.1093/jb/mvv027

43. Honda S, Kagoshima M, Wanaka A, et al. Localization and functional coupling of HGF and c-Met/HGF receptor in rat brain: implication as neurotrophic factor. Brain Res Mol Brain Res 1995;32(2):197–210, doi:10.1016/0169-328x(95)00075-4

44. Rehman R, Miller M, Krishnamurthy SS, et al. Met/HGFR triggers detrimental reactive microglia in TBI. Cell Rep 2022;41(13):111867, doi:10.1016/j.celrep.2022.111867

