## Supplemental Table for "Hepatocyte Growth Factor/MET Activator Rescues Working Memory Deficits After Repeated Mild Traumatic Brain Injury"

**Supplemental Table 1**: Animal weights prior to each repeated surgery.

| | | 3x Sham | | | | | 3x mTBI | | | | |
|---|---|---|---|---|---|---|---|---|---|---|---|
| Animal Sex | Surgery | N | Mean Weight (g) | Weight Std Err (g) | Weight Max (g) | Weight Min (g) | N | Mean Weight (g) | Weight Std Err (g) | Weight Max (g) | Weight Min (g) |
| F | Surgery 1 | 5 | 166.0 | 6.0 | 185 | 154 | 14 | 188.6 | 7.6 | 233 | 150 |
| F | Surgery 2 | 5 | ***187.2*** | 7.0 | 213 | 173 | 14 | ***202.9*** | 7.2 | 238 | 139 |
| F | Surgery 3 | 5 | ***190.6*** | 4.9 | 209 | 181 | 14 | ***208.7*** | 5.8 | 240 | 165 |
| M | Surgery 1 | 7 | ***199.4*** | 8.8 | 237 | 174 | 26 | ***205.7*** | 7.2 | 296 | 160 |
| M | Surgery 2 | 7 | ***238.6*** | 9.3 | 276 | 208 | 26 | ***235.3*** | 7.7 | 320 | 148 |
| M | Surgery 3 | 7 | ***249.4*** | 9.8 | 284 | 223 | 26 | ***250.6*** | 7.1 | 326 | 200 |

Red: Comparison to females; **Bold**: comparison to Surgery 1

**Supplemental Table 2**: Animal righing reflex during each repeated surgery.

| | | 3x Sham | | 3x mTBI | |
|---|---|---|---|---|---|
| Animal Sex | Surgery | Mean Righting Reflex (mm:ss.0) | Mean Righting Reflex Std Err (mm:ss.0) | Mean Righting Reflex (mm:ss.0) | Mean Righting Reflex Std Err (mm:ss.0) |
| F | Surgery 1 | 03:31.2 | 00:24.8 | 06:05.2 | 00:48.4 |
| F | Surgery 2 | 02:02.8 | 00:09.8 | 04:53.6 | 00:40.7 |
| F | Surgery 3 | 02:59.8 | 01:04.3 | 04:20.9 | 00:30.8 |
| M | Surgery 1 | 04:47.0 | 00:45.1 | 07:02.8 | 00:27.8 |
| M | Surgery 2 | 02:38.7 | 00:14.4 | 05:31.5 | 00:20.1 |
| M | Surgery 3 | 02:28.7 | 00:33.2 | 04:26.7 | 00:22.0 |
| Both | Surgery 1 | 04:15.4 | 00:24.8 | ***06:42.7*** | 00:29.5 |
| Both | Surgery 2 | 02:23.8 | 00:19.2 | ***05:18.2*** | 00:10.5 |
| Both | Surgery 3 | 02:41.7 | 00:17.7 | ***04:24.6*** | 00:31.6 |

Red: Comparison to 3x Sham; **Bold**: comparison to Surgery 1
