## Supplemental Figure 1 for "Hepatocyte Growth Factor/MET Activator Rescues Working Memory Deficits After Repeated Mild Traumatic Brain Injury"

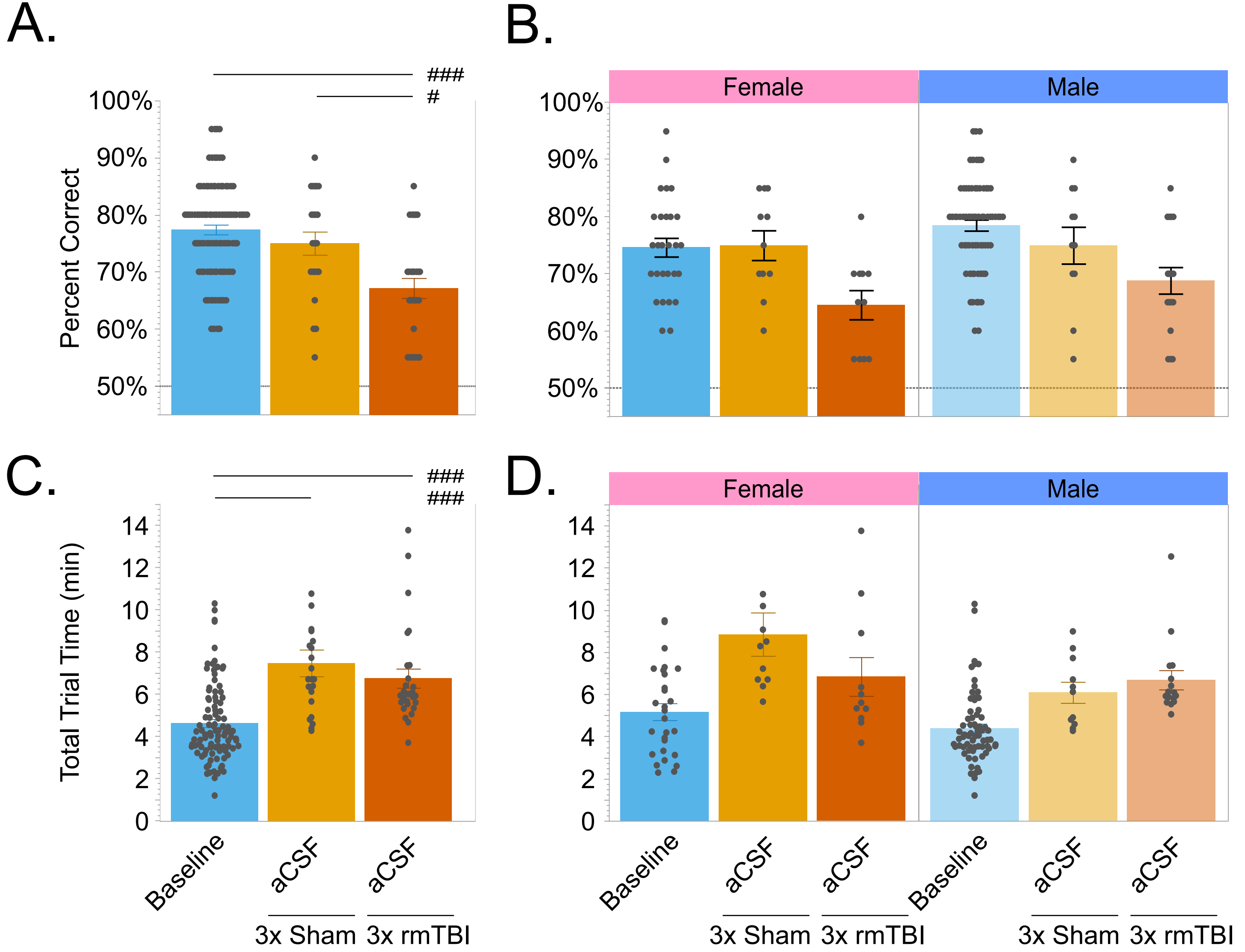


**Supplementary Figure 1.** Performance in the T-maze delayed alternation task of working memory during Baseline and after repeated sham or mTBI surgeries. 20 trials; 50% is chance performance. A) Performance in the T-maze task for all subjects and (B) separated out by sex. C) Total duration of T-maze testing for all subjects and (D) separated out by sex. #p<0.05, ##p<0.01, ###p<0.001, Tukey's HSD.
