## Supplementary File for "Hepatocyte Growth Factor/MET Activator Rescues Working Memory Deficits After Repeated Mild Traumatic Brain Injury"

Supplementary Material: **Detailed Materials and Methods**

**Experimental model and study participant details**

Animals

The male and female Long-Evans rats used in this study were obtained from both Inotiv (HsdBlu:LE #140; Inotiv Inc. West Lafayette, IN) and Charles River (Crl:LE #006; Charles River Laboratories, Raleigh, NC). Rats were single-housed with environmental enrichment in an AAALAC accredited temperature- and humidity-controlled environment on a 12/12-hour reverse light cycle. Housing under reverse light cycle ensured mild TBI surgeries and behavioral testing occurred during the animal’s active period. Access to food was regulated (5-10% bodyweight of food/day) during behavioral training and testing to maintain animals at 85–100% free-feeding weight and motivated to perform the behavioral task. All procedures were in accordance with NIH guidelines and approved by the Rowan University Institutional Animal Care and use Committee.

Cell Lines

Madin-Darby canine kidney (MDCK) cells were obtained from American Type Culture Collection (ATCC; NBL-2; #CCL-34).

**Experimental Models**

Mild Traumatic Brain Injuries

Repeated mild TBIs were performed using the electronic cortical contusion injury device (cCCI Model 6.3; Custom Design and Fabrication, Inc. Sandston, VA). Immediately after removal from isoflurane anesthesia (2-3% in 95%O2:5%CO2), animals were placed prone on a Marmarou foam block^1,2^ under the impactor (5 mm tungsten hemispherical tip). The skull, cleared of skin and periosteum, was impacted 2.5 mm caudal to bregma (5.5 m/s, 2.5mm depth, 100ms dwell). The animal was unrestrained and resting on the foam block which allowed the head to accelerate and move unrestrained following impact.

T-Maze Behavioral Test

The T-maze (90 cm wide × 65 cm long; corridor dim 10 cm wide × 10 cm high) was constructed from clear polycarbonate mounted on a white table^3-7^. The walls of the testing suite were lined with white plastic to minimize distal visual cues. Masking white noise was generated from a speaker 0.5 m behind the center of the maze measuring 60 db at the intersection of the ‘T’. The maze was wiped with 40% ethanol to minimize olfactory cues between each session^8^. Once learned, performance in this task may improve over multiple training/testing sessions. To maintain stable performance within a 75-85% correct range while waiting for surgery, delay duration during training was increased in 5-s increments when performance exceeded 90%. This approach ensured all animals learned the “rule” of the maze (i.e. delayed non-match to location) and is widely used to assess pharmacological manipulations on working memory performance^3,6,9-12^. Furthermore, our approach is functionally similar to non-human training protocols for primate tests of working memory^10^.

**Method Details**

Reagents

Dihexa (MedChemExpress Inc.; 1.0 pmol or 1.0 nmol/2 uL) was dissolved in 2.5% dimethyl sulfoxide (**DMSO**) and Dulbecco's Phosphate Buffered Saline (**DPBS**; #02–0117; VWR ). The HGF/MET antagonist, Hinge (**KDYIRN**, CHI Scientific; 300 pmol/2 μL) was dissolved in DPBS.

Bupivicane (2.5 mg/kg SC, #061842; Covetrus Inc.; Portland, Maine, US) and Rimadyl (4.0 mg/kg SC, #083911; Covetrus Inc.) were used as a surgical local anesthetic and post-operative analgesic.

Dulbecco’s modified Eagle’s medium (**DMEM**; #16777-124; VWR ) with fetal bovine serum (10%; #76419-586; VWR ) were used to grow MDCK cell colonies. Hepatocyte growth factor/scatter factor (HGF/SF; #HY-P73101; MedChemExpress) was used as a positive control for MDCK cell migration. E-Cadherin antibodies (#3195T; Cell Signaling Technology Inc.) and the secondary antibody Alexaflour 488 (#A21206; Invitrogen/ThermoFischer Inc.) was used to visualize CDMK cells.

T-Maze Task of Working Memory

Animals were allowed to acclimate to the vivarium for up to a week before starting training in the T-maze task of working memory. Training and testing in this delayed-nonmatch to position task were similar to that previously described^3,12,13^. Animals were trained to enter the T-maze goal arm opposite the last one visited to obtain food reward (1/8 froot loop, 24 mg; delivered by hand). Animals that reached a performance criteria of 80% ±5% accuracy over 21 trials (0 seconds delay, 1 session/day) in 3 consecutive sessions were randomly assigned to undergo repeated sham or closed-head mTBI surgeries and drug treatment. The first trial of the T-maze task reflects a spontaneous response arm choice by the animals and is not analyzed as a working memory trial. Stable performance at performance criteria for 3 days was used as an animal’s baseline performance measure. Under food regulation, animals reached stable performance criterion after approximately 40-50 sessions (20 trials/session, 1 session/day, 800-1000 trials). Training individual animals to a performance criteria ensures that animals have learned the “rule” of the maze (i.e. delayed non-match to location) and mirrors training protocols for testing the effects of pharmacological manipulations of working memory and those used for primate tests of working memory^3,6,9-12^. To maintain stable performance within a 75-85% correct range while animals were waiting for surgery, delay duration during training was increased in 5-s increments when performance exceeded 90%.

Immediately after the last sham or closed-head mTBI surgery, food regulation was reinstated. Animals were assigned an inter-trial delay, 5-sec. longer than delays used for baseline and underwent testing daily (1-5 days post final surgery). Testing sessions were conducted identically to training sessions.

Repeated mild TBI surgeries

Animals received repeated sham or closed-head mTBI surgeries (n=3 over 1 week) after being trained in the T-maze working memory task. Briefly, the midline of the skull was exposed and periosteum removed from animals under isoflurane anesthesia (2–3% in 95%O2:5%CO2). Animals were removed from anesthesia, placed on a Marmarou foam block^1,2^ in the eCCI device. The impactor tip was placed touching the skull, midline, 2.5 mm caudal to bregma, retracted, and impacted the skull (5.5 m/s, 2.5mm depth, 100ms dwell). Animals were removed from the eCCI device, placed on a sterile field in the supine position, and the time to exhibit a righting reflex was recorded. Animals were re-anesthetized, the skull was examined for bone fracture and/or hematoma, and the incision closed. During the series of repeated sham or closed-head mTBI surgeries animals were placed on *ad lib* food access.

Cannula implantation

During the final sham/mTBI surgery, cannula (26 ga. RWD, Sugar Land, TX) were placed above the right lateral ventricle (unilateral right side; -1.0AP, -1.4ML, -1.4DV) after the righting reflex and re-anesthetization. Jeweler's screws (304 stainless steel, M1x3mm) and dental acrylic anchored the cannula to the skull. Screws did not puncture dura. Sham animals experienced identical repeated surgical procedures but did not receive mTBI. Immediately after the final surgery, food regulation was resumed (5-10% bodyweight of food/day).

Intracerebroventricular drug infusions

For intracerebroventricular (ICV) infusions, vehicle (DPBS; 2 μL**)** or dihexa (1.0 pmol or 1.0 nmol/2 uL) was loaded into a 30-gauge infusion needle (800-00288-01; RWD Inc.; 2.0 mm projection) attached to PE10 tubing and 5 ul microsyringe (#87919; Hamilton Co. Inc.). Vehicle or Dihexa was infused at a rate of 2 µL/min under control of a microprocessor equipped infusion pump (#AL-300; World Precision Instruments, Sarasota, FL) into unrestrained, awake animals. Needles remained in the ventricle for 30 seconds following infusions. ICV infusions of Hinge (300 pmol/2 μL) were performed as a separate infusion, 5 minutes prior to dihexa infusions. Animals were returned to their home cage between infusion and working memory testing. Animals were returned to their home cage for the remaining 5 min before working memory testing. This high dose of dihexa was chosen from prior studies demonstrating an improvement in memory performance^14,15^.

MDCK cells

MDCK cells were initially cultured at 37°C in a 5% CO2 in air using DMEM + fetal bovine serum (10%). Subcultures were grown in six-well plates to form isolated, confluent colonies. Colonies were washed twice with DPBS and media was replaced with serum-free DMEM. DPBS, dihexa (10-1000 pM), HGF (0.34 μM), or dihexa (1000 pM)+HGF (0.34 μM) was added to the media and plates were incubated at 37°C with 5% CO_2_ for 10 hours. Media was removed and cells were fixed with 3.7% formalin in phosphate buffered saline followed by immunofluorescent labeling for E-Cadherin (#3195T; Cell Signaling Technology Inc.) and Alexaflour 488 (#A21206; Invitrogen/ThermoFischer Inc.).

**Quantification and Statistical Analysis**

All analyses were performed within the mixed model framework of JMP (ver. 18.2.1, JMP Statistical Discovery LLC.) and included AnimalID as the random variable. Statistical significance was set at p < 0.05. Tukey’s HSD post- hoc test was used to evaluate the effect of Injury, Surgery #, and Sex for the analysis of animal weights and righting reflex times. Tukey’s HSD and Dunnett-Hsu post-hoc test was used to compare performance in the T-maze task across injury groups and to compare drug effects to baseline performance.

Baseline working memory performance (% correct over 20 trials) and session duration was averaged for 3 training sessions immediately prior to surgery. Baseline performance from each experimental group was combined for statistical analyses. Working memory performance after repeated sham or mTBI was averaged over days 3-5. Working memory performance on post-injury days 1-2 were excluded from analysis considering animals motivation was re-equilibrating after reinstatement of food regulation and prior studies indicate that the effects of cMET signaling enhancement are found after 2-3 days of treatment^14,15^. Animals and individual testing days were excluded from analysis resulting from: surgical issues, environmental stressor, experimenter error.

E-Cadherin labeling of MDCK cells was quantified by imaging cell cultures (Keyence BZ-X710 microscope) and manually segmenting out individual colonies into individual photomicrographs. Cellpose-SAM^16^ running in a Google Colaboratory environment as a Jupyter Notebook was used to segment and identify cell membranes prior to importing these into ImageJ (National Institutes of Health, USA). The identified membranes were then scaled by 80% to allow quantification of E-Cadherin labeling intensity within the cytoplasm. Statistical analysis was performed within the mixed model framework of JMP using Cell_ID as the random variable. A Dunnett-Hsu post-hoc test was used to compare individual drug effects to baseline performance. statistical significance was set at p < 0.05.

**Code availability**

Cellpose-SAM is available by the authors at <https://www.github.com/mouseland/cellpose>. Scripts for recreating the analyses and figures are available at <https://github.com/DevilbissLab/ManuscriptAnalyses>.

**References**

1.	Marmarou CR, Prieto R, Taya K, et al. Marmarou Weight Drop Injury Model. In: Animal Models of Acute Neurological Injuries. (Chen J, Xu ZC, Xu X-M, et al. eds.) Humana Press: Totowa, NJ; 2009; pp. 393-407.

2.	Marmarou A, Foda MA, van den Brink W, et al. A new model of diffuse brain injury in rats. Part I: Pathophysiology and biomechanics. J Neurosurg 1994;80(2):291-300, doi:10.3171/jns.1994.80.2.0291

3.	Berridge CW, Devilbiss DM, Andrzejewski ME, et al. Methylphenidate preferentially increases catecholamine neurotransmission within the prefrontal cortex at low doses that enhance cognitive function. Biol Psychiatry 2006;60(10):1111-1120, doi:10.1016/j.biopsych.2006.04.022

4.	Devilbiss DM, Berridge CW. Cognition-enhancing doses of methylphenidate preferentially increase prefrontal cortex neuronal responsiveness. BiolPsychiatry 2008;64(7):626-635

5.	Devilbiss DM, Jenison RL, Berridge CW. Stress-induced impairment of a working memory task: role of spiking rate and spiking history predicted discharge. PLoS Comput Biol 2012;8(9):e1002681, doi:10.1371/journal.pcbi.1002681

6.	Spencer RC, Devilbiss DM, Berridge CW. The cognition-enhancing effects of psychostimulants involve direct action in the prefrontal cortex. Biol Psychiatry 2015;77(11):940-50, doi:10.1016/j.biopsych.2014.09.013

7.	Berridge CW, Devilbiss DM, Martin AJ, et al. Stress degrades working memory-related frontostriatal circuit function. Cereb Cortex 2023;33(12):7857-7869, doi:10.1093/cercor/bhad084

8.	Dudchenko PA. How do animals actually solve the T maze? Behavioral neuroscience 2001;115(4):850-60

9.	Verma A, Moghaddam B. NMDA receptor antagonists impair prefrontal cortex function as assessed via spatial delayed alternation performance in rats: modulation by dopamine. JNeurosci 1996;16(1):373-379

10.	Arnsten AF. Stress impairs prefrontal cortical function in rats and monkeys: role of dopamine D1 and norepinephrine alpha-1 receptor mechanisms. ProgBrain Res 2000;126(183-192

11.	Murphy BL, Arnsten AF, Goldman-Rakic PS, et al. Increased dopamine turnover in the prefrontal cortex impairs spatial working memory performance in rats and monkeys. Proc Natl Acad Sci U S A 1996;93(3):1325-9

12.	Devilbiss DM, Spencer RC, Berridge CW. Stress Degrades Prefrontal Cortex Neuronal Coding of Goal-Directed Behavior. Cereb Cortex 2016, doi:10.1093/cercor/bhw140

13.	Horvat L, Foschini A, Grinias JP, et al. Repetitive mild traumatic brain injury impairs norepinephrine system function and psychostimulant responsivity. Brain Res 2024;1839(149040, doi:10.1016/j.brainres.2024.149040

14.	McCoy AT, Benoist CC, Wright JW, et al. Evaluation of metabolically stabilized angiotensin IV analogs as procognitive/antidementia agents. The Journal of pharmacology and experimental therapeutics 2013;344(1):141-54, doi:10.1124/jpet.112.199497

15.	Benoist CC, Kawas LH, Zhu M, et al. The procognitive and synaptogenic effects of angiotensin IV-derived peptides are dependent on activation of the hepatocyte growth factor/c-met system. The Journal of pharmacology and experimental therapeutics 2014;351(2):390-402, doi:10.1124/jpet.114.218735

16.	Pachitariu M, Rariden M, Stringer C. Cellpose-SAM: superhuman generalization for cellular segmentation. bioRxiv 2025;2025.04.28.651001, doi:10.1101/2025.04.28.651001
